# Endangered Species Act listing is linked with greater research effort for U.S. butterflies

**DOI:** 10.64898/2026.05.30.729000

**Authors:** Rachel L. Walsh, Nich W. Martin, Ivone de Bem Oliveira, Jaret C. Daniels, Robert P. Guralnick, Akito Y. Kawahara

## Abstract

Conservation strategies for at-risk species can be aided significantly by research on topics such as ecology, life history, and threats, yet research effort is lacking for many species facing elevated extinction risk. Here we investigated whether listing under the U.S. Endangered Species Act (ESA) was associated with research effort for U.S. butterflies, and whether that effort was higher before or after ESA listing. We found that ESA-listed species had significantly more peer-reviewed publications than non-listed species after accounting for species range and taxonomic family. Further, we showed that more papers were published per year after ESA listing than before. These findings confirm that ESA-listed species benefit from greater research attention that can support data-informed conservation efforts. However, the relative scarcity of studies prior to ESA listing, as well as the lack of research for many unlisted, at-risk taxa, underscores the need for proactive, strategic research effort to inform conservation action.

## Introduction

Insect declines are now widely documented across taxa and regions. Wagner et al. (2021) estimates global insect declines of 1-2% per year, and Edwards et al. (2025) estimates a 22% loss in total U.S. butterfly abundance from 2000-2020. However, data that can be used to inform conservation actions are lacking. Walsh & Figueroa (2026) found that 88.5% of described North American insect and arachnid species have unknown extinction risk levels. Only 2% of invertebrates–and just 1.3% of insects–have been assessed for extinction risk by the International Union for the Conservation of Nature (IUCN), compared to 84% of vertebrates (*Summary Statistics*, 2025). Donaldson et al. (2017) found that invertebrates had significantly lower conservation research effort than vertebrates across geographic regions. Although substantial gaps remain in our understanding of insect declines, the documented losses are alarming given the critical roles that insects play in ecosystems. There are millions of insect species (Stork, 2018), many of which perform valuable ecosystem services that contribute hundreds of billions of dollars annually to the global economy (Losey & Vaughan, 2006). As E.O. Wilson aptly noted, invertebrates are “the little things that run the world” (Wilson, 1987).

Scientific data are critical for designing effective conservation plans and making decisions that support the strategic allocation of limited resources (Gerber et al., 2018; Joseph et al., 2009), but research effort is unevenly distributed and lacking for many at-risk species (Caldwell et al., 2024; Donaldson et al., 2017; Martín-López et al., 2009; Troudet et al., 2017). Studies have shown biases in research effort linked to various traits including body size, perceived charisma, range size, phylogeny, and human socioeconomic factors within species’ ranges (Brodie, 2009; Brooke et al., 2014; de Lima et al., 2011; Ducatez & Lefebvre, 2014; Guedes et al., 2023; Martín-López et al., 2009; Ridley et al., 2024; Santos et al., 2020). However, other authors have found no consistent relationship between IUCN designations of “Data Deficient” or “Critically Endangered” and research effort for invertebrates (Jarić et al., 2017). Funding patterns are also skewed: 82.9% of conservation funding is directed towards vertebrates, leaving the majority of at-risk taxa without support (Guénard et al., 2025). Addressing these imbalances requires systematic efforts to identify gaps and strategically direct research and conservation resources to areas of high impact, especially toward understudied groups, particularly insects.

The U.S. Endangered Species Act (ESA) is the primary legal mechanism for protecting at-risk species and their habitats in the United States (Greenwald et al., 2019; Walsh & Figueroa, 2026). ESA listing provides safeguards against take (i.e., to harass, harm, hunt, trap or kill), mandates the development of species recovery plans, and requires designation of critical habitat (Endangered Species Act, 1973). It is often assumed that listing also contributes to greater research attention and resource allocation. However, this assumption has rarely been tested empirically, and the relationship between ESA listing and research effort remains poorly understood (Weijerman et al., 2014). If ESA listing is associated with increased research activity, the resulting knowledge gains could strengthen conservation planning and recovery action for many at-risk species (Martín-López et al., 2009). Conversely, a lack of association could highlight a disconnect between legal protection and the scientific information needed to support recovery, underscoring the need for additional incentives to promote research.

In this study, we examined research effort allocation among U.S. butterflies. We focused on butterflies because they are among the best-studied insect groups and are a prominent locus of conservation initiatives, making them a model for assessing research trends within insects. We tested two hypotheses: 1) ESA listing is associated with greater overall research effort, and 2) publication rates increase following species listing. Our goal was to evaluate a case study for how legal protection is linked with scientific effort, in order to better understand allocation of research attention broadly.

## Materials and Methods

### Data collection

We obtained a list of all butterfly species and subspecies that occur in the United States from NatureServe Explorer in July 2025 (NatureServe Explorer, 2025). We chose to include subspecies because several butterfly subspecies are listed under the ESA. From NatureServe, we also obtained synonyms, ESA statuses, and species distributions in terms of occurrence within U.S. states and Canadian provinces and territories. Taxa that had an ESA status of “Endangered” or “Threatened” were categorized as “Listed,” and we assigned a category of “Not Listed” to all others. We generated a coarse proxy for species range size by counting U.S. states and Canadian provinces or territories in which each taxon had a state- or province-level NatureServe ranking.

We obtained taxonomic family assignments from GBIF and LepTraits, prioritizing the latter where discrepancies occurred (GBIF, 2025; Shirey et al., 2022). One family name (Hesperiidae for *Telegonus*) was manually assigned because it was not found in either database. We cross-validated ESA statuses from NatureServe with the USFWS ECOS database (USFWS, 2025) and accordingly classified Myrtle’s Silverspot (*Argynnis zerene myrtleae*) and Bay Region Checkerspot (*Euphydryas editha bayensi*) as “Listed.” Because the NatureServe dataset excludes species solely from U.S. territories, our analysis is limited to butterflies found within U.S. states.

We conducted systematic literature searches through the Web of Science Starter API accessed using *rwosstarter* (Casajus, 2023; Clarivate Developer Portal, 2025). The search results represent a systematic sample that is not comprehensive but representative of peer-reviewed publications for U.S. butterflies. For each taxon in our dataset, we searched the scientific name and synonyms (see supplementary materials). We retained all publications retrieved, so that the criterion for inclusion was that the scientific name appeared in the text. We did not filter by subject area due to API limitations and the large number of records. By retaining all publications retrieved for each taxon, our approach is conservative: any association between ESA listing and research effort must emerge against a broad and heterogeneous body of literature extending beyond conservation-focused studies. For each taxon, we exported all retrieved records and combined them into a single dataset, retaining the search term used to retrieve each paper.

To test our hypothesis that ESA listing is associated with greater research effort, we calculated the all-time total number of papers retrieved per taxon as the response variable. Bibliometric records included a unique identifier and publication year for each paper, allowing us to also calculate the number of publications per taxon per year. Annual publication counts served as the response variable for testing our second hypothesis that publication rates increase following ESA listing. We obtained ESA listing years for each listed butterfly from the USFWS ECOS database and used these dates to categorize publications occurring before or after listing. Papers published the same year as listing were conservatively categorized as “Before Listing,” under the assumption that funding and research activities preceded formal designation. Bibliometric data were downloaded in July 2025, and ESA listing dates were obtained in October 2025 (Casajus, 2023; Clarivate Developer Portal, 2025, NatureServe Explorer, 2025; USFWS, 2025).

### Data analysis

To test our first hypothesis that ESA listing is associated with greater research effort for U.S. butterflies, we used the total number of publications retrieved per taxon across all years as the response variable. The main predictor was ESA Status (“Listed” versus “Not Listed”). We included taxonomic family and a coarse estimate of range size as fixed effects, with range serving as a proxy for geographic exposure. We fit models using *glmmTMB* (McGillycuddy et al., 2025), selecting a zero-inflated negative binomial model to reflect the structure of overdispersed count data with excess zeros. All three predictors (ESA Status, Family, and Range) were included in both the count and zero-inflation submodels. We included predictors in both submodels because we hypothesized that our predictors would be associated with research attention expressed both as publication counts (count submodel) and odds of being completely unstudied (zero-inflation submodel). For both hypotheses, we evaluated model performance using AIC, pseudo-R^2^, and residual diagnostics, and variance inflation factors (VIF) where appropriate, and selected the specified model as the best performing. For each analysis, we calculated estimated marginal means using *emmeans* (Lenth & Piaskowski, 2017), and implemented residual checks with *DHARMa* (Hartig, 2016).

Our second hypothesis was that publication rates increase following ESA listing. The response variable was the number of papers published per year for each taxon, and the main predictor was publication timing relative to ESA listing (“Before” versus “After”). We fit models using *glmmTMB* (McGillycuddy et al., 2025) and found that a negative binomial model was the best performing given overdispersed count data. To control for annual variation in overall research activity, we included an offset equal to the natural logarithm of one plus the total number of papers for all U.S. butterfly species in each year. We also included a species-level random intercept to account for repeated observations (before and after) for each taxon. Coarse range estimate and taxonomic family were not included in this model because they are time-invariant species attributes accounted for by the random intercept. We did not include a listing year term because the model estimates differences in annual rather than cumulative paper counts, so there is no need to account for accumulated papers after listing for an early listing year, and vice versa.

## Results

Our dataset included 1,436 U.S. butterfly species and subspecies, of which 33 are listed under the U.S. Endangered Species Act (7 Threatened; 26 Endangered). Our systematic literature searches retrieved 4,531 unique papers (1909-2025) for 453 taxa (Table S1).

Consistent with our first hypothesis, the full model detected significantly greater research effort for ESA-listed taxa relative to unlisted taxa after accounting for range and family (pairwise contrast: *p* = 0.012, effect = 12.426, 95% CI = 2.770 – 22.082, Table S2). Model predictions indicate that ESA-listed species are associated with an average of 14.8 papers, compared with

2.4 papers for unlisted species (Figure 1, Table 1). In the count submodel, predicted means were 12.8 papers for listed species and 2.5 papers for unlisted species (Table S3). Estimates from the zero-inflation submodel were highly uncertain; therefore, we refrain from drawing conclusions regarding the influence of ESA status on the structural zero process (Table S4). This is likely due to the extremely low odds of being completely unstudied for listed species.

**Table 1.** Estimated marginal means from the full model for total number of papers by ESA status (H1) and for annual publication rate per unit of research exposure by timing of ESA listing (H2).

| Estimated Mean Papers by ESA Status (Full Model) |  |  |  |  |
| --- | --- | --- | --- | --- |
| ESA Status | Estimate | SE | LCL | UCL |
| Listed | 14.790 | 4.934 | 5.120 | 24.460 |
| Not Listed | 2.364 | 0.247 | 1.880 | 2.848 |

| Estimated Mean Papers per Year by Timing of Listing |  |  |  |  |
| --- | --- | --- | --- | --- |
| Timing | Estimate | SE | LCL | UCL |
| Before Listing | 0.00027 | 0.00010 | 0.00007 | 0.00046 |
| After Listing | 0.00100 | 0.00033 | 0.00034 | 0.00165 |

| Estimated Mean Papers by ESA Status (Full Model) |  |  |  |  |
| --- | --- | --- | --- | --- |
| ESA Status | Estimate | SE | LCL | UCL |
| Listed | 14.790 | 4.934 | 5.120 | 24.460 |
| Not Listed | 2.364 | 0.247 | 1.880 | 2.848 |
| Estimated Mean Papers per Year by Timing of Listing |  |  |  |  |
| Timing | Estimate | SE | LCL | UCL |
| Before Listing | 0.00027 | 0.00010 | 0.00007 | 0.00046 |
| After Listing | 0.00100 | 0.00033 | 0.00034 | 0.00165 |

**Figure 1.**
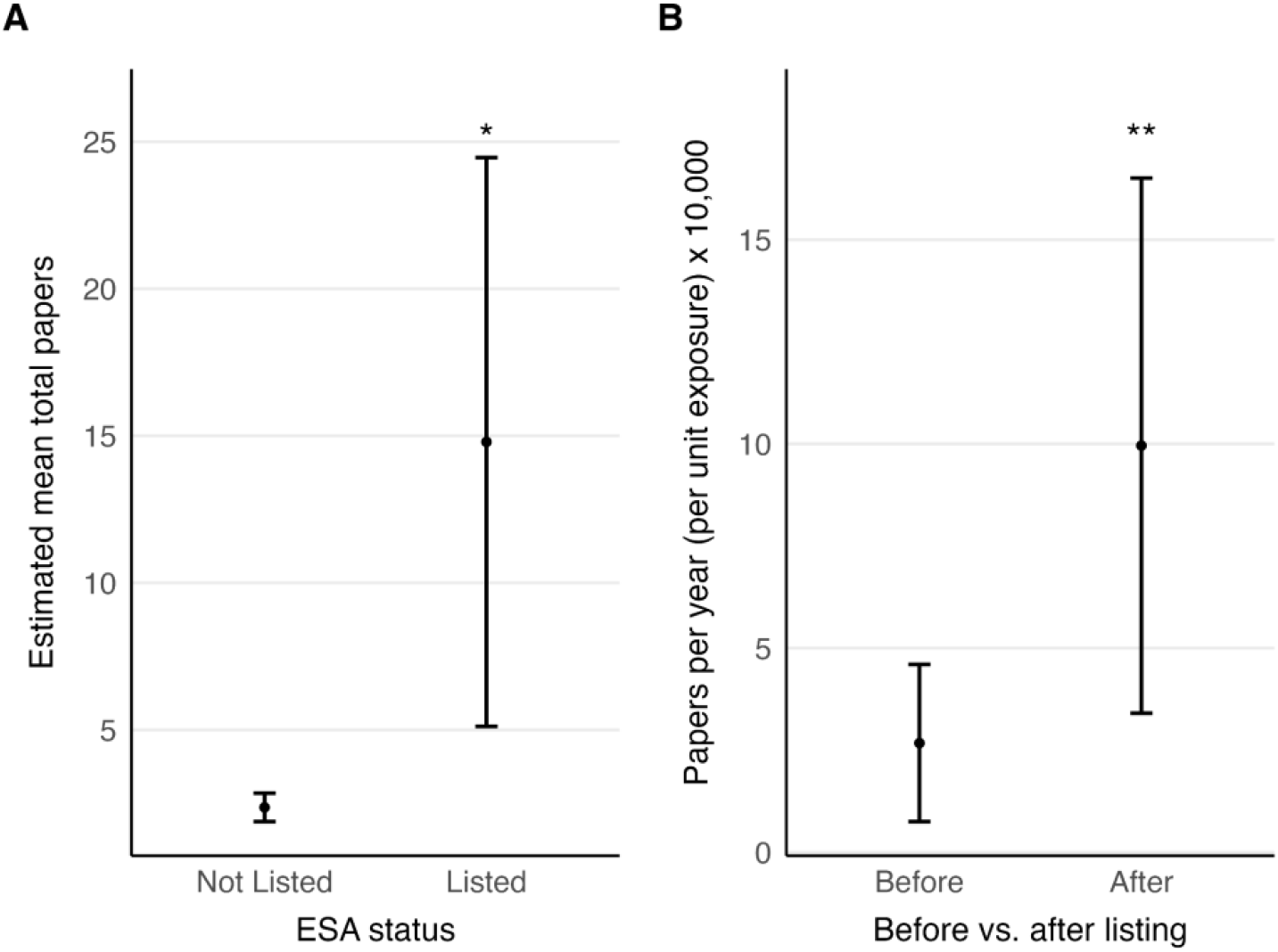
A) Model-estimated mean total papers from the full model for ESA-listed versus unlisted taxa (H1). A zero-inflated negative binomial model estimated an unconditional mean of 14.8 papers for listed taxa, and 2.4 papers for unlisted taxa (*p* = 0.012, effect = 12.426, 95% CI = 2.770 – 22.082). B) Model-estimated mean annual publication rate before and after ESA listing per unit of exposure multiplied by 10,000. A negative binomial model estimated a pre-listing publication rate of 0.0003 and a post-listing rate of 0.0010 per unit of research exposure, or a 3.7x higher publication rate after listing (*p* = 0.004, effect = 0.0007, 95% CI = 0.0002 – 0.0012).

Estimated marginal means from the full model, evaluated across representative values of coarse range size (0, 20, 40, and 60 states or provinces), revealed a consistent increase in research effort with increasing range, with all pairwise contrasts showing significance (Figure 2, Table S5, Table S6). For taxonomic family, Papilionidae had significantly more papers than the grand mean across families (*p* = 0.033, effect = 5.877, 95% CI = 0.851 – 10.903), whereas Hesperiidae (*p* = 0.002, effect = -2.075, 95% CI = -3.269 – -0.882), Lycaenidae (*p* = 0.002, effect = -2.156, 95% CI = -3.343 – -0.970), and Riodinidae (*p* = 0.016, effect = -2.212, 95% CI = -3.843 – - 0.581) had significantly fewer (Figure 3, Table S8). Pieridae showed a nonsignificant positive deviation from the grand mean (*p* = 0.495, effect = 0.792, 95% CI = -1.103 – 2.687), and Nymphalidae did not differ significantly (*p* = 0.725, effect = -0.225, 95% CI = -1.480 – 1.030). Nagelkerke’s pseudo-R^2^ indicates moderate explanatory power (pseudo-R^2^ = 0.31).

**Figure 2.**
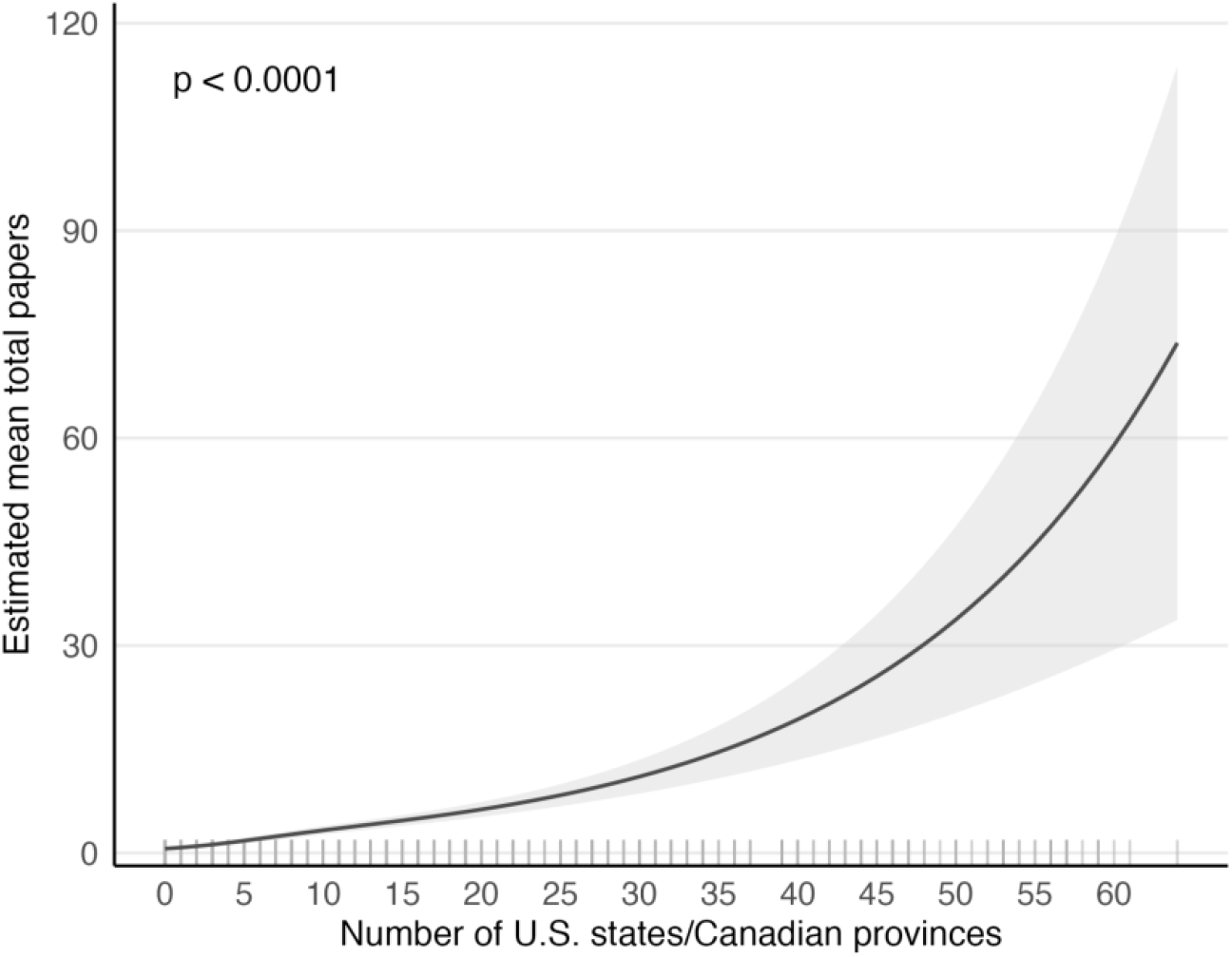
Relationship between coarse range estimate (number of U.S. states and Canadian provinces/territories) and estimated mean total papers from the full zero-inflated negative binomial model (conditional submodel: *p* < 0.0001, IRR = 1.057, 95% CI = 1.046 – 1.069).

**Figure 3.**
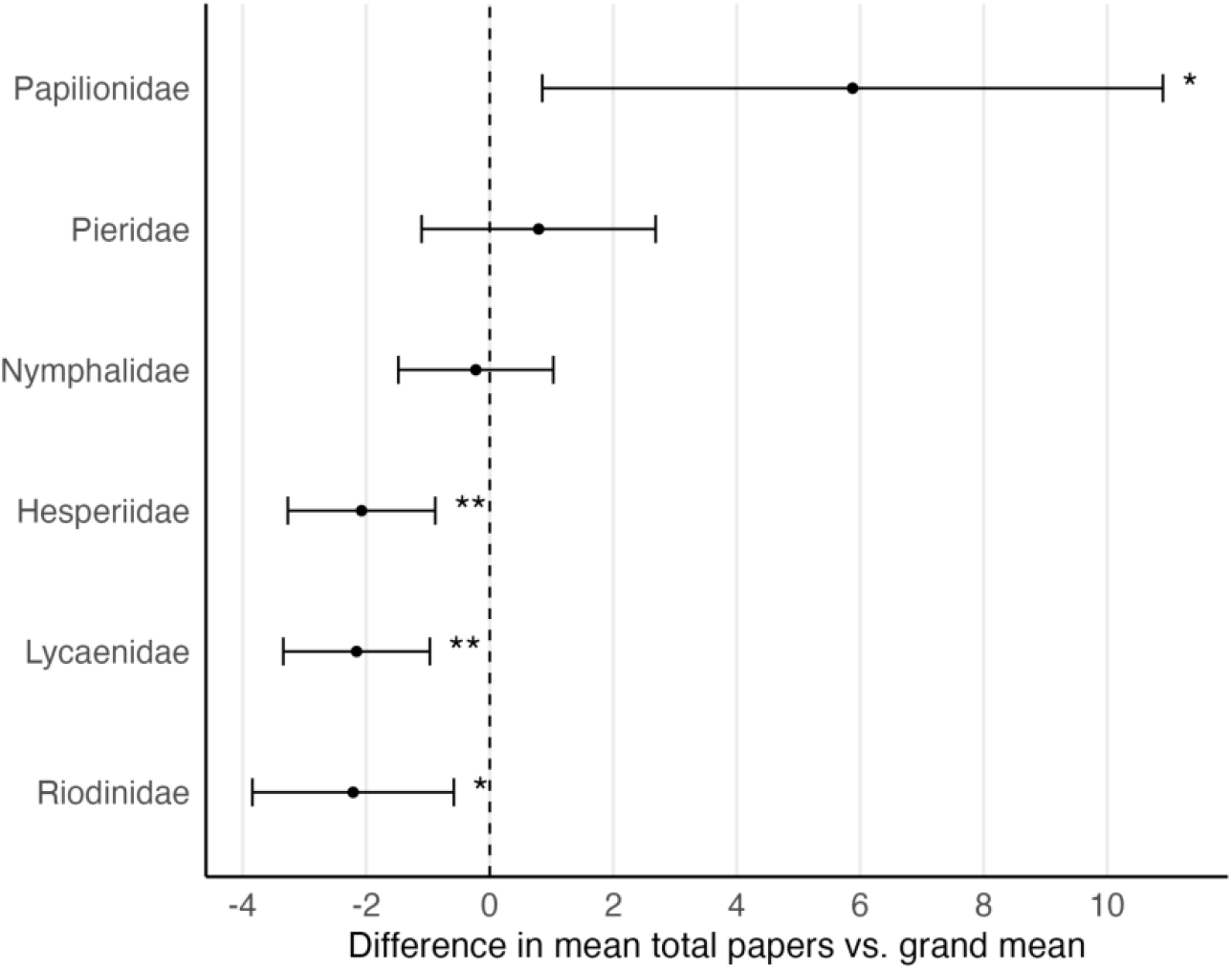
Difference between estimated mean total papers and the grand mean across families for each of the six taxonomic families. The full zero-inflated negative binomial model estimated significantly more papers than the grand mean for Papilionidae (*p* = 0.033, effect = 5.877, 95% CI = 0.851 – 10.903) and significantly less for Hesperiidae (*p* = 0.002, effect = -2.075, 95% CI = -3.269 – -0.882), Lycaenidae (*p* = 0.002, effect = -2.156, 95% CI = -3.343 – -0.970), and Riodinidae (*p* = 0.016, effect = -2.212, 95% CI = -3.843 – -0.581), and a non-significant difference from the grand mean for Pieridae and Nymphalidae.

Consistent with our second hypothesis, annual publication rates were higher after ESA listing than before (pairwise contrast: *p* = 0.004, effect = 0.0007, 95% CI = 0.0002 – 0.0012). The model estimated an approximately 3.7-fold increase in publication rate following listing after accounting for annual publication exposure (Figure 1, Table S9). Based on Nagelkerke’s pseudoR^2^, the model showed moderate explanatory power (pseudo-R^2^ = 0.39).

## Discussion

Our results showed that ESA-listed taxa were linked to substantially greater research effort than unlisted taxa, even after accounting for range size and family-level effects. Publication rates also increased significantly after ESA listing. These findings indicate that ESA designation is linked with significantly enhanced scientific knowledge production for at-risk U.S. butterfly species.

Research effort was positively associated with geographic range. Taxa occurring in a higher number of states or provinces received greater scientific attention, likely due to increased exposure and accessibility (Brooke et al., 2014; Ducatez & Lefebvre, 2014; Guedes et al., 2023). We also detected significant family-level differences, suggesting that taxonomic identity captures unmeasured traits that influence research investment, such as detectability, body size, charisma, historical research emphasis, or applied relevance (Bellon, 2019; Brooke et al., 2014; Guedes et al., 2023). Papilionidae (swallowtails), for example, received significantly greater attention, potentially reflecting their larger size and conspicuous appearance. In contrast, Hesperiidae, Lycaenidae, and Riodinidae were associated with lower publication effort, perhaps due to their smaller size or less visually striking phenotypes. Research attention for Pieridae was above average, which might reflect the fact that several species in this family are agricultural pests, leading to greater research output for this group.

In practice, research effort is intertwined with ESA listing through status assessments and recovery planning (Endangered Species Act, 1973). The statute requires that listing decisions must be made based on the “best scientific data available” and mandates recovery plans and periodic status reviews for listed taxa (Endangered Species Act, 1973), processes that can engage the scientific community and stimulate research production. Scientific research outputs are therefore both a prerequisite for listing and a consequence of it (Martín-López et al., 2009).

Limitations of the Web of Science Starter API prevented topic-level classification of publications. Furthermore, we did not quantitatively assess additional factors such as body size or pest status. Additionally, our study focused on peer-reviewed literature because it provides a systematically indexed and readily accessible record of research output. We suggest that future studies incorporate grey literature, evaluate additional predictors, and categorize papers by subject to better understand drivers of research allocation and their implications for conservation.

Our findings present important takeaways for conservation research and practice. Scientific knowledge strengthens conservation efforts by informing allocation of limited resources (Gerber et al., 2018; Joseph et al., 2009). Research can help identify priority habitats (Noon, B. R., & McKelvey, K. S., 1996), delineate movement corridors (Kautz et al., 2006), identify drivers of decline (Shirey & Ries, 2023), and determine which life stages are most vulnerable (Heppell, 1998). Thus, greater research attention for ESA-listed species may enhance conservation strategies for species known to be in need of protection.

Although butterflies are one of the best-studied insect groups, 69% of U.S. taxa had zero publications in our systematic search. In addition to unlisted species and species with smaller geographic ranges, families with smaller and less conspicuous species (Hesperiidae, Lycaenidae, Riodinidae) received significantly less attention. These results suggest that the broader insect fauna, which lacks comparable attention and is dominated by small, cryptic taxa, likely experiences even deeper research gaps. Despite mounting evidence of widespread diversity losses, only 4% of insect species–representing more than 60% of described animal diversity–are protected under the ESA (Stork, 2018; Zhang, 2011). While demonstrating greater research attention for ESA-listed species, our results also imply that this elevated attention likely does not reach most declining species. Therefore, we highlight a mismatch between the scale of insect declines and the distribution of legal protection and scientific investment, leaving most at-risk taxa outside institutional mechanisms that mobilize action.

Significant challenges remain, but targeted actions could reduce disparities in research allocation. Greater coordination among academic institutions, government agencies, and non-governmental organizations is needed to align research priorities with conservation needs and speed assessment processes (Caldwell et al., 2024; Guénard et al., 2025; Walsh & Figueroa, 2026). These efforts can strengthen the scientific foundation for effective conservation action.

### Conclusion

This study demonstrates that research effort for U.S. butterflies is strongly associated with ESA listing status. Listed taxa receive substantially greater scientific attention and experience higher publication rates following designation. Although we do not infer causality, our results indicate that formal conservation status is closely linked to the concentration of scientific effort. At the same time, widespread butterfly and insect declines occur largely outside the scope of ESA protection, and many at-risk taxa appear not to benefit from the elevated research attention associated with listing. A central contribution of this work is demonstrating that scientific investment remains uneven relative to the scale of biodiversity loss, even within one of the best-studied insect groups. Addressing ongoing insect declines will require not only sustained support for ESA-listed species, but also proactive strategies to generate ecological knowledge for a wide range of vulnerable taxa. More broadly, our findings highlight the need for systematic, evidence-based approaches for prioritizing conservation research in ways that better align scientific effort with extinction risk under limited resource conditions.

## Funding Statement

Funding for this project was provided by the School of Natural Resources and Environment, Institute of Food and Agricultural Sciences, and the Florida Museum of Natural History at the University of Florida. The authors declare that they have no conflicts of interest relevant to this manuscript.

### Data Availability Statement

All data and code used in this study are publicly available via Zenodo. Data are available at: https://doi.org/10.5281/zenodo.19499930

Scripts are available at: https://doi.org/10.5281/zenodo.19499842

## Supporting Information

Additional supporting information may be found in the online version of this article:

## Appendix S1: Supplementary tables and figures

## Notes

https://doi.org/10.5281/zenodo.19499930

https://doi.org/10.5281/zenodo.19499842

### Competing Interest Statement

The authors have declared no competing interest.

