## Appendix S1: Supplementary tables and figures for "Endangered Species Act listing is linked with greater research effort for U.S. butterflies"

**Table S1:** Observed counts of taxa with and without papers by ESA listing status from a systematic literature search for 1,436 U.S. butterflies in July 2025.

| Taxa With and Without Published Papers by ESA Status | | | | |
|---|---|---|---|---|
| **ESA Status** | **Paper Group** | **Count (n)** | **Total** | **Percent (%)** |
| Listed | Has Papers | 25 | 33 | 75.76 |
| Listed | Zero Papers | 8 | 33 | 24.24 |
| Not Listed | Has Papers | 428 | 1403 | 30.51 |
| Not Listed | Zero Papers | 975 | 1403 | 69.49 |

**Table S2:** Pairwise contrast between listed and unlisted taxa showing a difference in means of 12.4 total papers from the full zero-inflated negative binomial model ($p = 0.012$).

| Difference in Mean Paper Counts by ESA Status | | | | | |
|---|---|---|---|---|---|
| **Contrast** | **Estimate** | **SE** | **LCL** | **UCL** | **P-value** |
| Listed - Not Listed | 12.426 | 4.926 | 2.770 | 22.082 | 0.012 |

**Table S3:** Estimated marginal means for total number of papers by ESA status (count submodel).

| Estimated Mean Papers by ESA Status (Count Submodel) | | | | |
|---|---|---|---|---|
| **ESA Status** | **Estimate** | **SE** | **LCL** | **UCL** |
| Listed | 12.753 | 4.176 | 6.713 | 24.228 |
| Not Listed | 2.537 | 0.242 | 2.105 | 3.059 |

**Table S4:** Estimated probability of structural zero by ESA status (zero-inflation submodel).

| Estimated Probability of Structural Zero by ESA Status | | | | |
|---|---|---|---|---|
| **ESA Status** | **Pr(Structural Zero)** | **SE** | **LCL** | **UCL** |
| Listed | 0.000 | 0.000 | 0.000 | 1.000 |
| Not Listed | 0.192 | 0.063 | 0.097 | 0.344 |

**Table S5:** Estimated marginal means for total number of papers by coarse range estimate (full model).

| Estimated Mean Papers by Number of States (Full Model) | | | | |
|---|---|---|---|---|
| States/Provinces | Estimate | SE | LCL | UCL |
| 0 | 0.618 | 0.105 | 0.412 | 0.825 |
| 20 | 6.300 | 0.553 | 5.216 | 7.384 |
| 40 | 19.310 | 2.994 | 13.441 | 25.179 |
| 60 | 58.990 | 15.098 | 29.400 | 88.581 |

**Table S6:** Pairwise contrasts between selected values of coarse range estimate (*n* = 0, 20, 40, 60 states/provinces) showing a significant increase in mean total paper count with occurrence in an increasing number of states/provinces, after accounting for ESA status and family (full model).

| Difference in Mean Paper Counts by Number of States/Provinces | | | | | |
|---|---|---|---|---|---|
| Contrast | Estimate | SE | LCL | UCL | P-value |
| 0 - 20 | −5.681 | 0.515 | −6.690 | −4.673 | 0.000 |
| 0 - 40 | −18.691 | 2.995 | −24.561 | −12.821 | 0.000 |
| 0 - 60 | −58.372 | 15.113 | −87.993 | −28.751 | 0.001 |
| 20 - 40 | −13.010 | 2.632 | −18.169 | −7.850 | 0.000 |
| 20 - 60 | −52.690 | 14.822 | −81.741 | −23.640 | 0.002 |
| 40 - 60 | −39.681 | 12.217 | −63.626 | −15.736 | 0.006 |

**Table S7:** Estimated marginal means for total number of papers by taxonomic family (full model).

| Estimated Mean Papers by Family (Full Model) | | | | |
|---|---|---|---|---|
| Family | Estimate | SE | LCL | UCL |
| Hesperiidae | 1.526 | 0.220 | 1.095 | 1.958 |
| Lycaenidae | 1.445 | 0.219 | 1.016 | 1.874 |
| Nymphalidae | 3.376 | 0.486 | 2.424 | 4.329 |
| Papilionidae | 9.479 | 3.101 | 3.401 | 15.557 |
| Pieridae | 4.394 | 1.025 | 2.384 | 6.403 |
| Riodinidae | 1.389 | 0.735 | −0.050 | 2.829 |

**Table S8:** Main effect contrasts between taxonomic family and the grand mean across families showing a significantly higher mean paper count than the grand mean for Papilionidae ($p$ = 0.033), and a significantly lower mean paper count for Hesperiidae ($p$ = 0.002), Lycaenidae ($p$ = 0.002), and Riodinidae ($p$ = 0.016), after accounting for ESA status and family (full model).

Difference in Mean Paper Counts by Family vs. Grand Mean

| Contrast | Estimate | SE | LCL | UCL | P-value |
|---|---|---|---|---|---|
| Hesperiidae effect | −2.075 | 0.609 | −3.269 | −0.882 | 0.002 |
| Lycaenidae effect | −2.156 | 0.605 | −3.343 | −0.970 | 0.002 |
| Nymphalidae effect | −0.225 | 0.640 | −1.480 | 1.030 | 0.725 |
| Papilionidae effect | 5.877 | 2.564 | 0.851 | 10.903 | 0.033 |
| Pieridae effect | 0.792 | 0.967 | −1.103 | 2.687 | 0.495 |
| Riodinidae effect | −2.212 | 0.832 | −3.843 | −0.581 | 0.016 |

**Table S9:** Pairwise contrast between before and after ESA listing showing a mean difference of 0.0007 fewer papers per species per year before ESA listing at unit exposure ($p$ = 0.0044).

Difference in Mean Paper Counts by Timing of Listing

| Contrast | Estimate | SE | LCL | UCL | P-value |
|---|---|---|---|---|---|
| Before - After Listing | −0.0007 | 0.0003 | −0.0012 | −0.0002 | 0.0044 |

**Figure S1:** Observed total number of papers from a systematic literature search in July 2025 for all 33 ESA-listed U.S. butterfly taxa; bars are filled by taxonomic family.

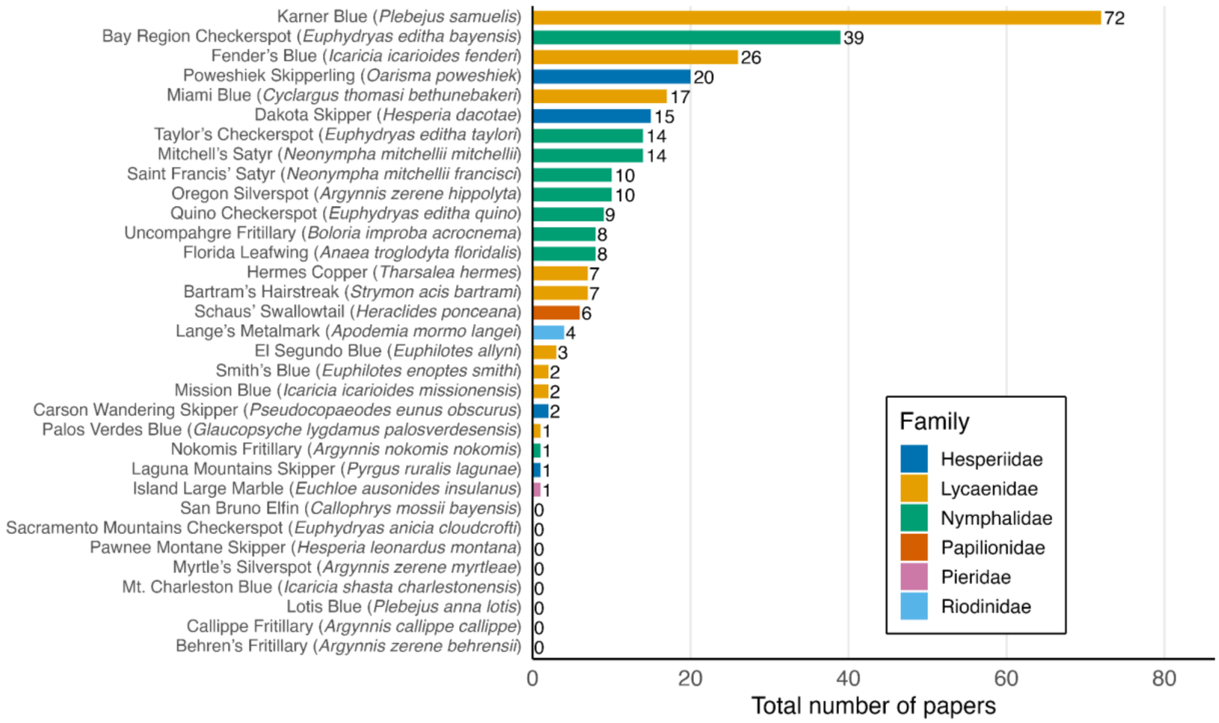


**Figure S2:** Observed total number of papers per species by taxonomic family from a systematic literature search in July 2025 for 1,436 U.S. butterfly taxa; points are filled by ESA status.

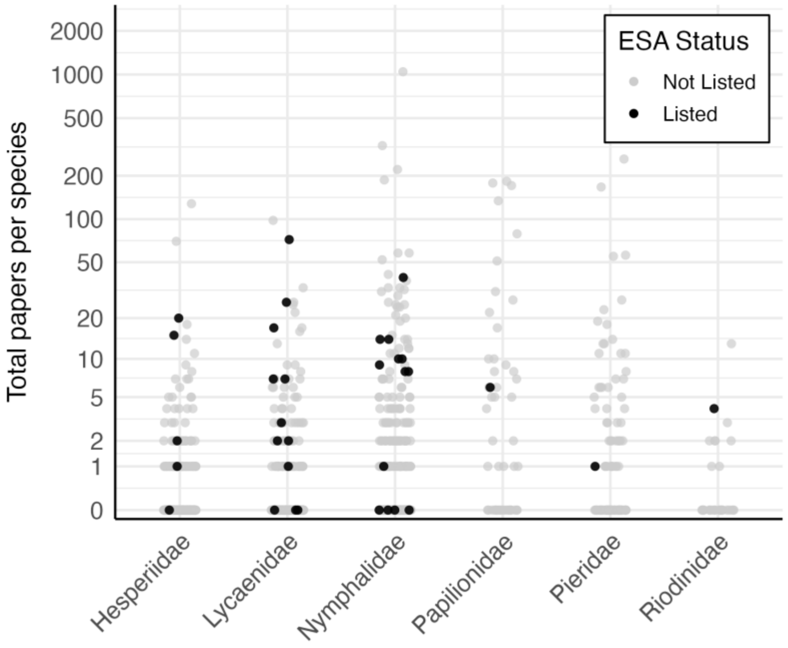


**Figure S3:** Observed number of papers per species per year by timing of ESA listing relative to publication year (before vs. after listing) from a systematic literature search conducted in July 2025 for all 33 ESA-listed U.S. butterfly taxa.

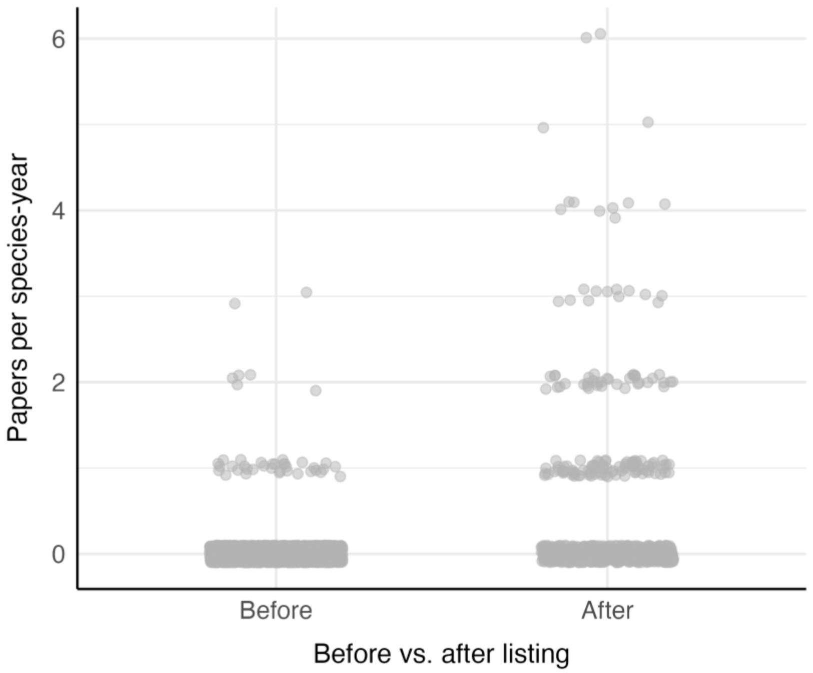


**Figure S4:** Observed number of papers per species per year over time filled by ESA status.

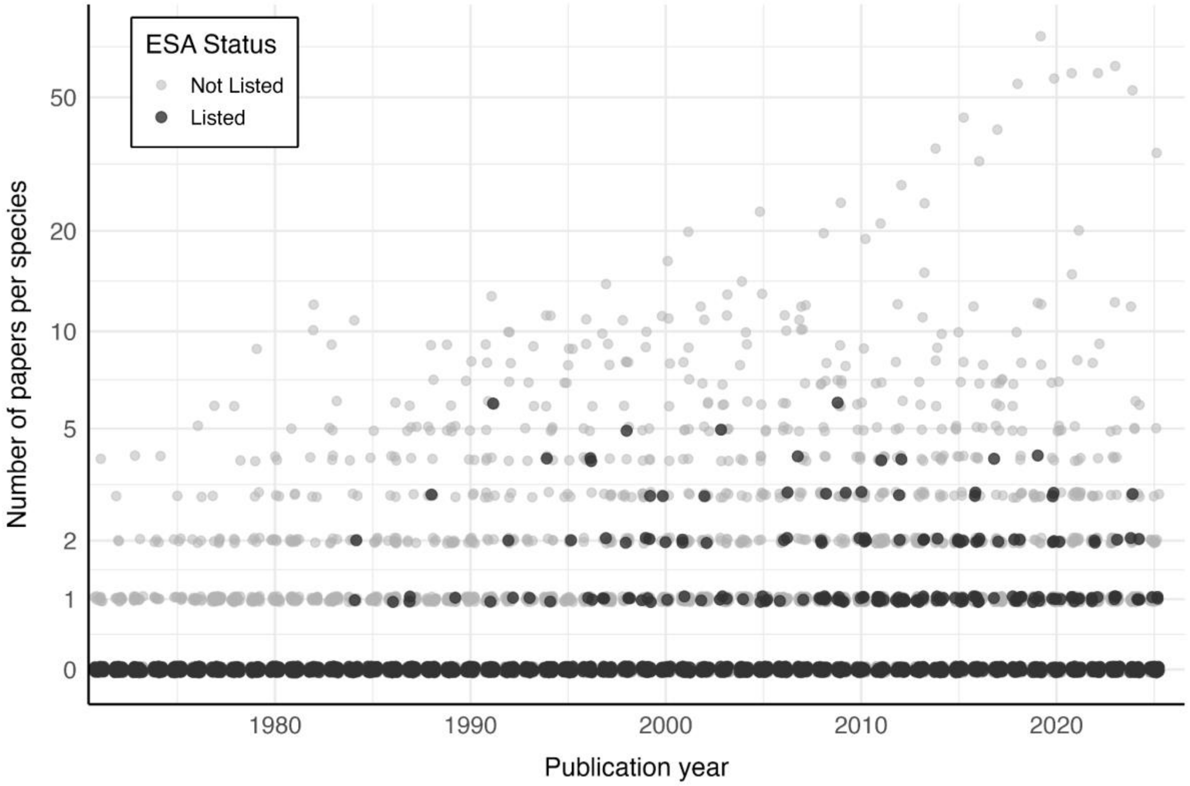


Data are available at: https://doi.org/10.5281/zenodo.19499930
Scripts are available at: https://doi.org/10.5281/zenodo.19499842
